# Ramping cells in rodent mPFC encode time to past and future events via real Laplace transform

**DOI:** 10.1101/2024.02.13.580170

**Authors:** Rui Cao, Ian M. Bright, Marc W. Howard

**Affiliations:** Department of Psychological and Brain Sciences, Boston University

## Abstract

In interval reproduction tasks, animals must remember the event starting the interval and anticipate the time of the planned response to terminate the interval. The interval reproduction task thus allows for studying both memory for the past and anticipation of the future. We analyzed previously published recordings from rodent mPFC (Henke et al., 2021) during an interval reproduction task and identified two cell groups by modeling their temporal receptive fields using hierarchical Bayesian models. The firing in the “past cells” group peaked at the start of the interval and relaxed exponentially back to baseline. The firing in the “future cells” group increased exponentially and peaked right before the planned action at the end of the interval. Contrary to the previous assumption that timing information in the brain has one or two time scales for a given interval, we found strong evidence for a continuous distribution of the exponential rate constants for both past and future cell populations. The real Laplace transformation of time predicts exponential firing with a continuous distribution of rate constants across the population. Therefore, the firing pattern of the past cells can be identified with the Laplace transform of time since the past event while the firing pattern of the future cells can be identified with the Laplace transform of time until the planned future event.

---

Imagine singing a favorite song. In the pause before the next line of lyrics, the memory of the previous line gradually recedes, while the planned time to sing the next line approaches. As time progresses, the precise moment to begin singing the next line becomes increasingly clear, culminating at the moment of vocalization. The ability to prepare and execute an action at a specific moment often requires us to track the flow of time internally, which is explicitly tested in interval production tasks (Jazayeri & Shadlen, 2015; Mauk & Buonomano, 2004; Merchant, Zarco, Pérez, Prado, & Bartolo, 2011; Narayanan, 2016). In a typical interval reproduction experiment, subjects experience a target duration and must later reproduce the duration through planned actions (e.g. Fig 1a). Just like the pause between two lines in a song, each moment throughout the reproduction phase is defined relative to a beginning point at *t* = 0 that recedes into the past as *t* increases and an endpoint at *t* = *T* . At each moment after *t* = 0 the endpoint is a distance *T−t* from the present, approaching from the future (see Fig 1b).

**Figure 1.**
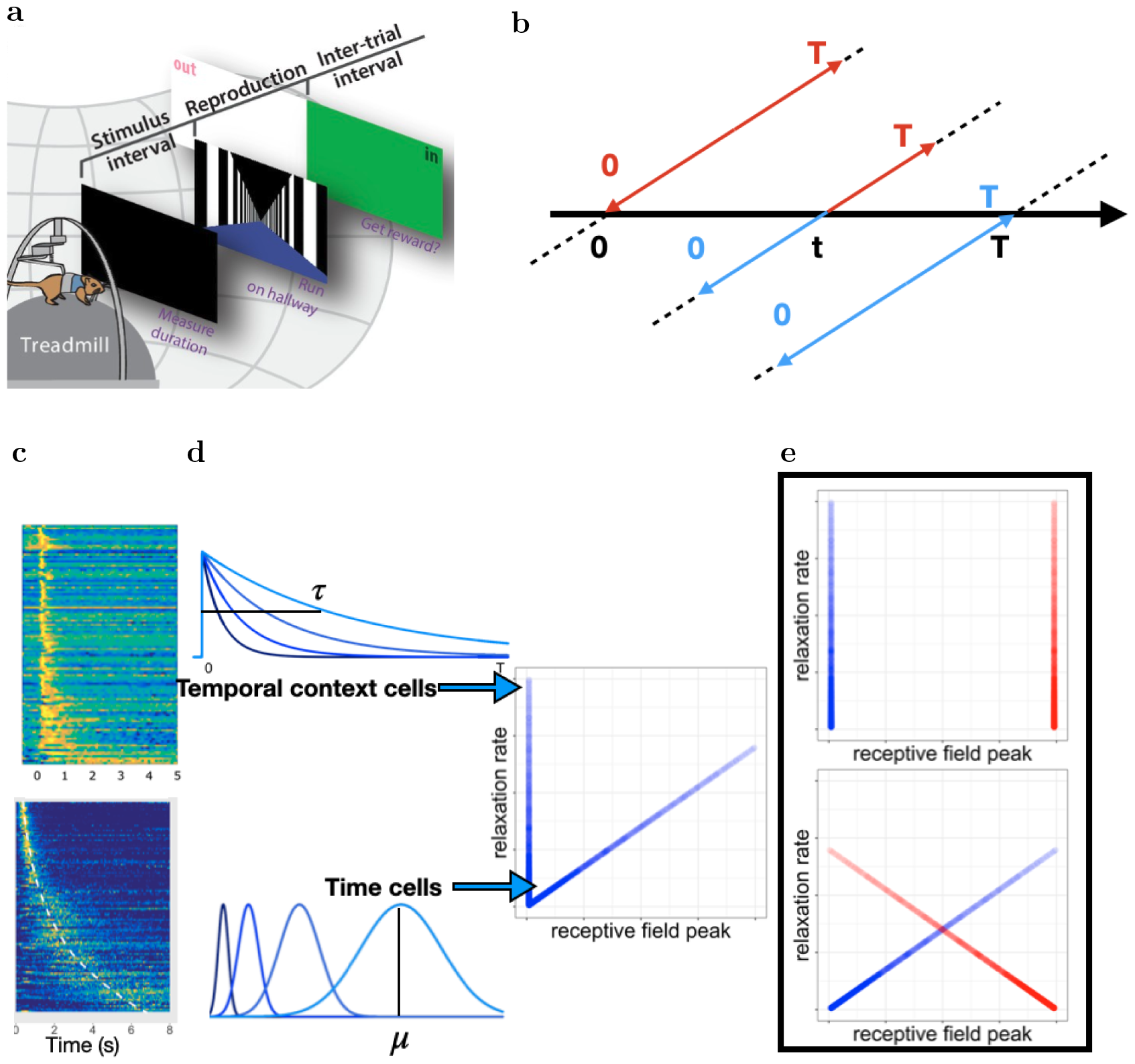
Neural representations of time elapsed since the beginning point and time remaining until the endpoint in interval reproduction tasks. **a**. Experimental task from Henke, et al., (2021). On each trial, a gerbil experiences a target stimulus duration, and then reproduces the duration by running on a treadmill. Here we limit our analysis to the neural activity during the reproduction phase (adapted from Henke, et al., 2021). **b**. Representation of the target interval in the reproduction phase. The horizontal line represents objective time, and the diagonal lines represent the target interval, which consists of time lapsed (*t*) since the beginning of the interval (blue) and time remaining (*T− t*, red). The beginning of the interval is in the past while the end of the interval is in the future during the reproduction phase. **c-d**. Time elapsed can be encoded by temporal context cells and time cells. **c**. Top panel shows previously reported temporal context cells, adapted from Bright, et al., 2020; the bottom panel shows time cells, adapted from Cao, et al., 2022. In both panels, each row illustrates the normalized firing rate of a cell where yellow indicates high firing rates and blue indicates low firing rates. **d**. Illustration of stimulated temporal context cells (top) and time cells (bottom). Time cells fire sequentially with widening receptive fields after an inciting event, resulting in a wide range of peak locations across cells. By contrast, a group of temporal context cells would reach their peak firing rate shortly after an event and relax back to baseline firing at different rates for different cells. Therefore, when receptive field peaks are plotted as a function of the relaxation rates in the left panel, temporal context cells form a narrow vertical stripe. **e**. Hypothetical properties of the neural populations during the reproduction phase with either temporal-context-cell-like representations (top) or time-cell-like representations (bottom). The temporal-context-cell-like representation would result in two vertical lines, one at the beginning and one at the end of the interval. The time-cell-like presentation would result in a wide range of receptive field peaks. The illustrations are presented symmetrically for easy visualization. The neural data is analyzed separately for the beginning and the end of an interval.

A number of hypotheses have been proposed to describe how we keep track of the time of future planned actions internally during tasks such as interval reproduction. Scalar expectancy theory (SET) proposed that time elapsed is tracked through an accumulator that counts the pulses generated by an internal clock after the beginning point (Gibbon, Church, Meck, et al., 1984). While models like SET primarily focus on behavioral data, later models incorporated neural data observed during timing tasks. There is a large animal recording literature that reports ramping activity during timing tasks; individual neurons monotonically increase or decrease their activity as time progresses (Jazayeri & Shadlen, 2015; Henke et al., 2021; Kim, Ghim, Lee, & Jung, 2013; Kunimatsu, Suzuki, Ohmae, & Tanaka, 2018; Mita, Mushiake, Shima, Matsuzaka, & Tanji, 2009; Narayanan, 2016; Xu, Zhang, Dan, & Poo, 2014). Therefore later theories implemented a single clock that accumulates the firing activities of a group of cells and reaches a threshold at the end of an interval (Merchant & Averbeck, 2017; Simen, Balci, de Souza, Cohen, & Holmes, 2011). More recently, it was proposed that the changing dynamics of neural population activity, in the form of sequentially activated cells (Bakhurin et al., 2017; Zhou, Masmanidis, & Buonomano, 2020) or neural trajectories in low-dimensional space (Remington, Narain, Hosseini, & Jazayeri, 2018; Henke et al., 2021), serves as an internal timing mechanism. Researchers applied recurrent neural networks (RNNs) to perform timing tasks (Laje & Buonomano, 2013; Remington, Egger, Narain, Wang, & Jazayeri, 2018) and implemented neural-inspired constraints to those RNNs (Cueva et al., 2020; Zhou, Masmanidis, & Buonomano, 2022) as a way to gain further insights into the population dynamics required for internal time tracking. Researchers specifically focused on restricting the dimension of RNNs by building low-rank RNNs and argued it led to improved generalization to untrained stimuli (Beiran, Meirhaeghe, Sohn, Jazayeri, & Ostojic, 2023).

These diverse models all share the underlying assumption that the neural representation changes with a few characteristic time scales across neurons for a given interval. SET model assumes a single poison process as the internal clock (Gibbon et al., 1984). Accumulator models typically assume that the neurons are noisy exemplars of the overall rate, and the ramping rate reflects aggregated firing rates across neurons (Simen et al., 2011). For low-rank RNN models, the dynamics^1^ have as many time constants as the rank of the RNN has eigenvalues (Jazayeri & Shadlen, 2015; Beiran et al., 2023). In all of those models, the typical time scale shared by the population only changes for different interval lengths.

Recent work on memory for the past has shown robust evidence that neural activity shows an effectively continuous set of time constants across neurons (Fig., 1c,d). “Time cells” fire sequentially during a delay following an event (MacDonald, Lepage, Eden, & Eichenbaum, 2011; Pastalkova, Itskov, Amarasingham, & Buzsáki, 2008). Time cells have been widely reported in the hippocampus (Cruzado, Tiganj, Brincat, Miller, & Howard, 2020; MacDonald et al., 2011; Mau et al., 2018; Pastalkova et al., 2008), but have also been reported in the medial entorhinal cortex (Kraus et al., 2015), the prefrontal cortex (Cruzado et al., 2020; Ning, Bladon, & Hasselmo, 2022; Tiganj, Cromer, Roy, Miller, & Howard, 2018), and striatum (Akhlaghpour et al., 2016; Mello, Soares, & Paton, 2015).

The peak times of time cells smoothly tile the delay interval; as the sequence advances time cells fire for progressively longer durations (Kraus, Robinson, White, Eichenbaum, & Hasselmo, 2013; Cao, Bladon, Charczynski, Hasselmo, & Howard, 2022). Time cells thus exhibit a continuous distribution of characteristic time constants. “Temporal context cells” (Fig. 1c top) also show a continuous distribution of time constants. Temporal context cells all respond shortly after some event, and relax their firing back to baseline at a variety of different rates (Bright et al., 2020; Tsao et al., 2018; Zuo et al., 2023). Rather than conveying information about how long ago an event happened via a continuous set of peak times (Fig. 1d bottom), temporal context cells show a continuity of relaxation time constants (Fig. 1d top).

The effectively continuous time constants in these two types of cells have similar properties. Within a time cell population, the density of peak times decreases linearly as time progresses, while the size of the corresponding temporal receptive windows linearly increases. Consequently the time of the triggering event is represented on a logarithmic time scale (Cao et al., 2022). The time constants of temporal context cells also overrepresent short time scales and have a smooth distribution with a long tail. Populations of temporal context cells are well-described *via* exponential relaxation, *e*^*−st*^ with a broad range of real *s* (Bright et al., 2020; Tsao et al., 2018). Thus one can identify the firing of a population of temporal context cells with the real Laplace transform of the time since the triggering event was experienced (Shankar & Howard, 2013).

To characterize the temporal information represented in a reproduction task using previously established methods for past time (Bright et al., 2020; Cao et al., 2022), we analyzed recently published recordings by Henke et al. (2021) from gerbil medial prefrontal cortex(mPFC) as they performed an interval production task (Fig.1a). During the measurement phase, the animals were presented with the target interval length *T*, sampled randomly from a set ranging from 3 - 7.5 s. During the reproduction phase they would time their running to match the target interval length. We restricted our analyses to the reproduction phase to investigate the temporal representation of the past action (beginning of the interval) and the future action (projected end of the interval). We observed separate groups of units that represent either time since the beginning of the delay or time until the end of the delay. These two groups contained qualitatively similar properties in their respective representations: both are characterized by a narrow range of peak values and a wide range of time constants, resembling the temporal context cells observed in Bright et al. (2020). Finally, we show that time in these groups is coded continuously, ruling out the possibility that time is represented via a small number of time scales.

## Results

A total of 1766 mPFC cells were recorded in Henke et al. (2021). We first identified cells with reliable temporal receptive fields using previously established methods (Bright et al., 2020). The method revealed two groups of temporal-responsive cells: cells with temporal receptive field preferentially peaking shortly after the start of the interval (331 cells see Fig. 2a) and cells with temporal receptive field preferentially peaking shortly before the end of the interval (391 cells, see Fig. 2b). We refer to these populations as past cells and future cells respectively. These two groups can also be appreciated from Fig. 3 in Henke et al. (2021). We proceed to analyze the 722 temporal responsive cells with hierarchical Bayesian models. We performed the same models and statistical tests on the remaining 1044 cells and found similar results (see Supplemental section 1).

**Figure 2.**
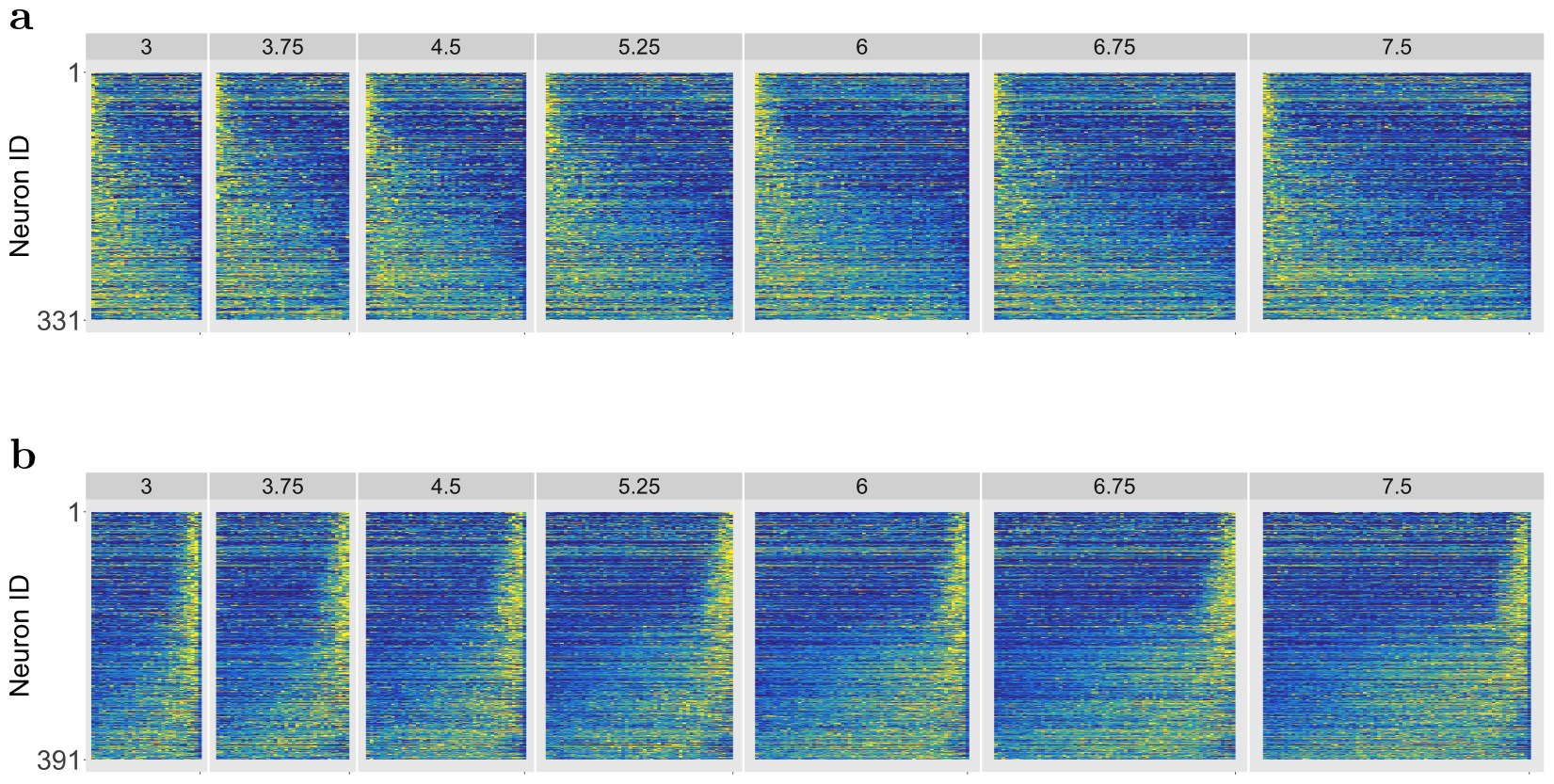
The start and the end of an interval are coded in two separate groups of cells across different intervals a. Heatmap of past cells during reproduction for each stimulus interval. Each row represents the normalized firing of a cell, linearly rescaled and averaged over all trials under the same stimulus length (marked for each column). Yellow indicates high firing and dark blue represents zero firing rate. Cells are sorted by the corresponding estimated time constant *τ* during the 5.25 s interval. **b**. Same as in **a**., but for the future cells.

**Figure 3.**
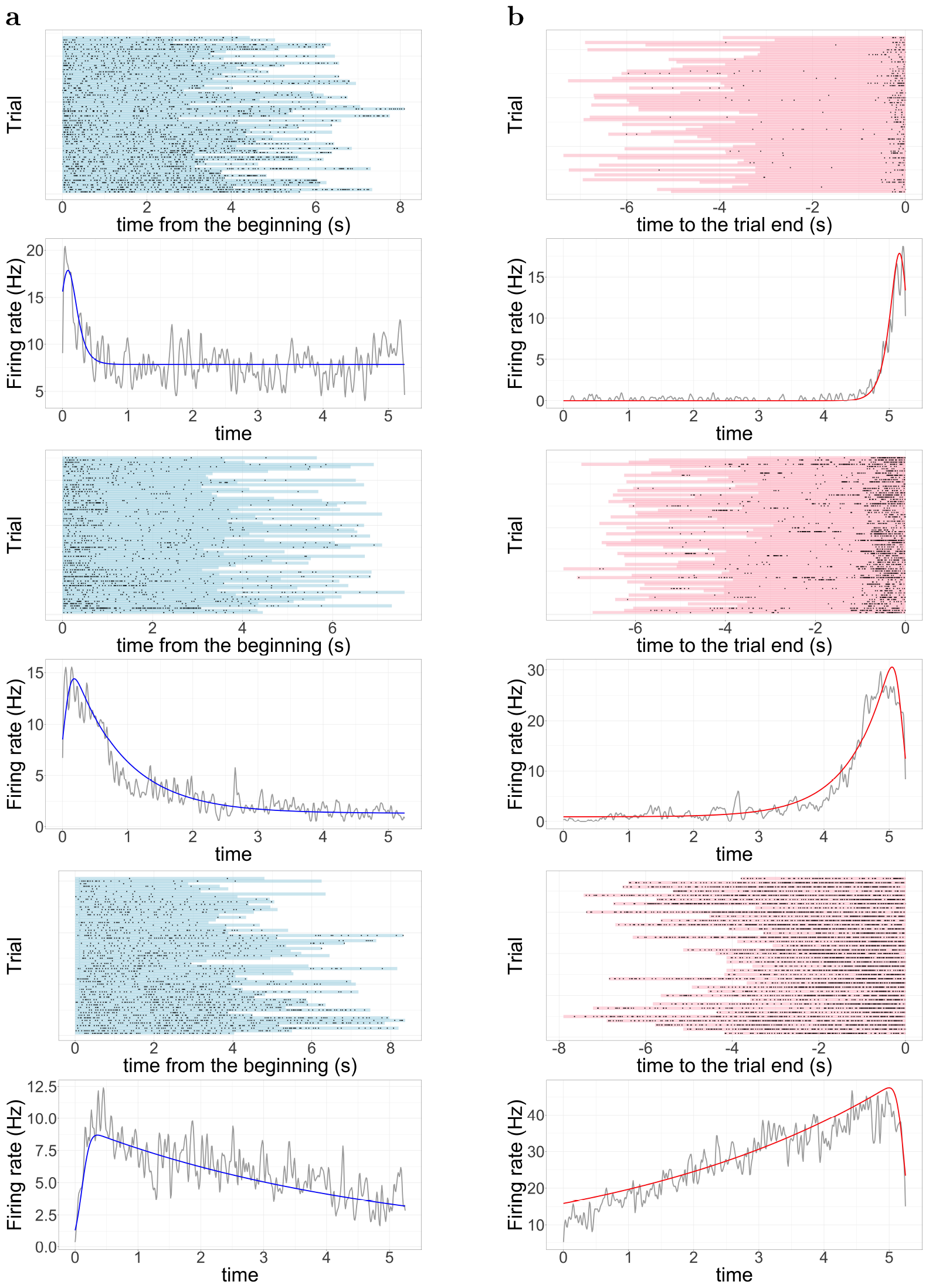
Example past cells (a) and future cells (b). Each figure contains three example cells with different time constants from the corresponding group. We plot each cell with the raster plot (top), and the averaged firing rate (bottom). In each raster plot, the dots represent the spike train and the length of color shade corresponds to the reproduced interval length of a given trial. For the past cells (a), the trials are aligned from the start of the interval; for the future cells (b), the trials are aligned from the end of the interval. In each average firing plot, the grey line represents the average firing rate of the cell after its activity for every trial linearly rescaled to a 5.25 s interval. The colored line represents the estimated temporal receptive field for the cell during a 5.25 s interval.

The hierarchical Bayesian model estimates temporal receptive fields with an exponentially modified Gaussian (ex-Gaussian) with two key parameters: 1) the peak parameter *µ* and 2) the time constant parameter *τ*, which indicates the time it takes for the cell to regress 63% back to its baseline firing. The ex-Gaussian receptive field is sufficiently expressive to allow observation of time cell populations and temporal context cell populations. Time cell populations would have a wide range of estimated peak locations *µ* as the sequence continuously tiles the delay (Fig. 1e bottom). Temporal context cell populations would have a narrow range of peak locations *µ* but a wide range of time constants *τ* that continuously tile the delay (Fig. 1e top).

To account for the changing temporal receptive fields for different interval lengths, we set the time constant *τ* for *T* = 5.25*s* as the baseline and assumed that the time constant for each trial, *τ*_*i*_ may change according to the reproduced interval length *T*_*i*_. Specifically, we assumed 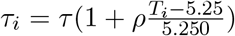 where *ρ* determines how much each cell rescales its temporal receptive field according to interval lengths. When *ρ* = 0, the cell codes for absolute time and does not change its temporal receptive field for different interval lengths while *ρ* = 1 means the cell linearly rescales its time constant for the interval lengths. Details about the model can be found in the method section. Consistent with previous analysis of the neural population (Henke et al., 2021), we also observed that cells largely rescaled to the interval length and reported the results in Supplemental section 3.

Example fits from the hierarchical Bayesian model can be found in Fig. 3 (for more examples see Fig. S2). For the past cells that preferentially peaked at the start of the interval, we estimated the temporal receptive field parameters since the start of the interval; as a function of time elapsed *t*. For the future cells that preferentially peaked at the end of the interval, we estimated the temporal receptive fields as a function of time until the end of the studied interval, *T − t*.

### The temporal receptive fields of mPFC cells are characterized by a narrow range of peak locations and a broad range of time constants

As described above, the temporal information within the interval can be coded through either a broad range of temporal receptive field peak locations *µ* (e.g. time cells), or a broad range of time constant values *τ* (e.g. temporal context cells). Therefore we focused our analysis on these two parameters for both past cells and future cells below.

Figure 4a summarizes the temporal response properties of the 331 past cells. To facilitate inspection of the overall firing pattern of each cell across *all* intervals, we linearly rescaled cell activities for each trial to 5.25 s and plotted the averaged activity for one cell in each row. The figure demonstrates that almost all the cells reached peak firing shortly after the start of an interval but different cells relax back to baseline firing at different rates within the interval. Figure 4c summarizes the temporal response properties of the 391 future cells. Almost all the future cells reached peak firing shortly before the end of the interval. Different cells ramped up their activity at different rates within the interval, mirroring the pattern for past cells.

**Figure 4.**
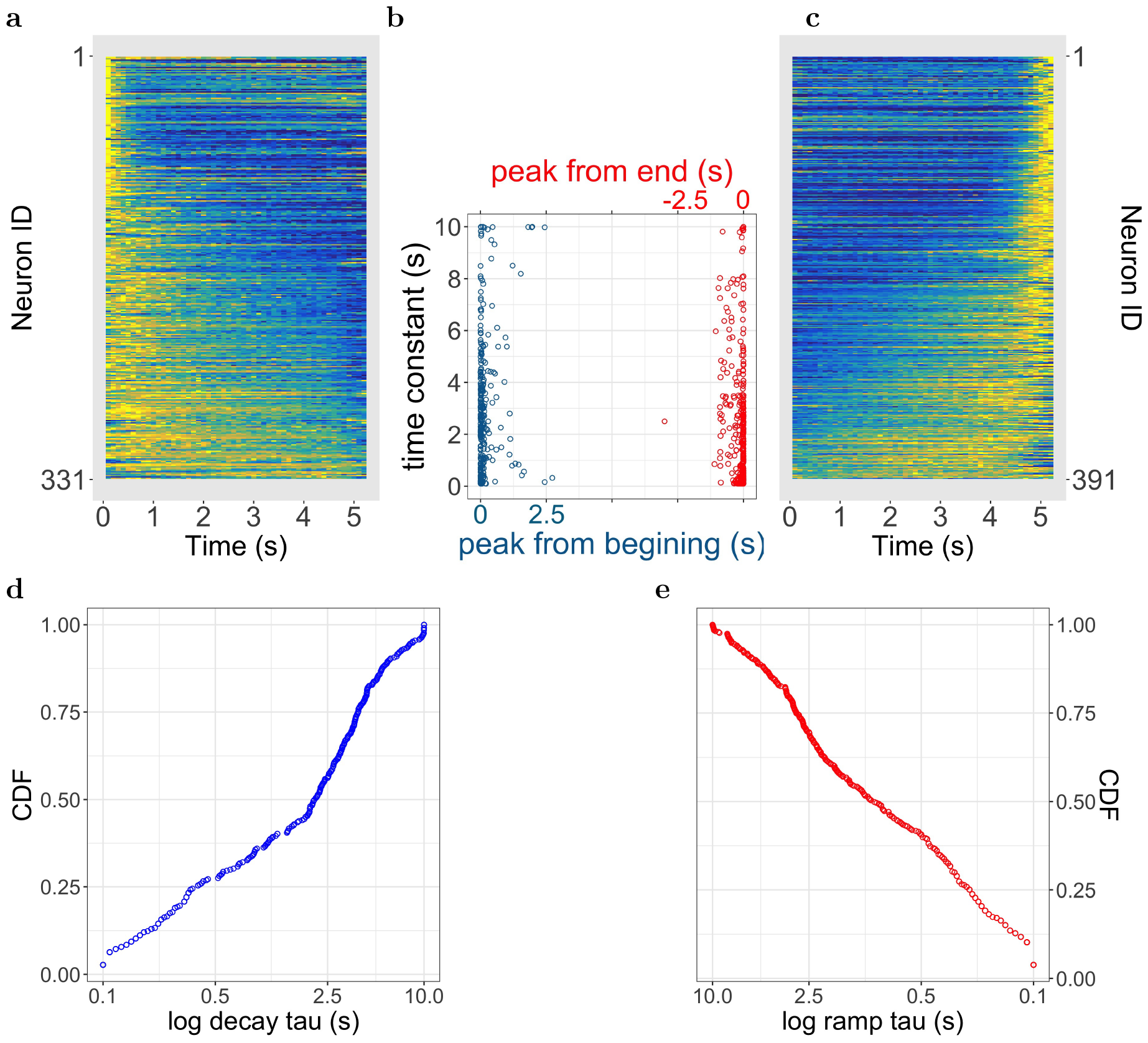
The start and end of interval is represented with continuous time constants. **a-c** mPFC cells tile the reproduced interval with a broad range of time constants from either the start or the end of the interval. **a**. Normalized firing rate of past cells. Cells are sorted based on time constant *τ* and plotted in the same way as figures in Fig. 2, except averaged over all the trials where every trial was linearly rescaled to 5.25 s. **b**. Receptive field peaks *µ* plotted as a function of estimated time constants *τ* for both past cells (blue) and future cells (red). For past cells, peak location *µ* was calculated from the start of the interval (blue) and for future cells, *µ* was calculated from the end of the interval (red). **c**. Normalized firing rate of future cells. Plotted in the same way as **a. d-e**. The cumulative density function (CDF) of time constants plotted on a log scale for past cells (d) and future cells (e). Each point represents a cell in the corresponding group.

Distributions of peak locations *µ* and time constants *τ* estimated through formal modeling of individual cells from the two groups confirmed these qualitative observations (see Fig. 4b). We reported and performed subsequent statistical analysis on the most frequent observations from the posterior distribution for simplification.

### The peaks of past cells cluster at the start of the interval

As can be appreciated from Fig. 4b, the estimated peak locations *µ* for past cells were distributed over a narrow range of small values (median = 0.01 s, interquartile range = 0 to 0.10 s, 90_*th*_ percentile = 0.41 s). Indeed most of the past cells reached peak firing rate shortly after the beginning of the interval (303/331 cells fire within 0.5 s). This pattern was true within each animal: for the 76 cells from animal 1, the median of estimated peak locations was 0.01 s with 90_*th*_ percentile = 0.39 s; for the 156 cells from animal 2, the median of estimated peak locations was 0.01 s with 90_*th*_ percentile = 0.23 s; for the 99 cells from animal 3, the median of estimated peak locations was 0.01 s with 90_*th*_ percentile = 1.11 s. This provides strong evidence against the hypothesis that time elapsed since the start of the interval is coded by a group of sequentially firing cells that tile the interval as one would expect for a population of time cells.

### The time constants of past cells are distributed over a broad range

Contrary to the narrow range for *µ*, the estimated time constants *τ* for the past cells resulted in a wide range of values (median = 2.11 s, interquartile range = 0.39 to 3.88 s, 90_*th*_ percentile = 6.11 s, see Fig. 4b). This pattern was true within each animal: for the 76 cells from animal 1, the median of estimated time constant was 1.94 s with interquartile range from 0.32 to 3.22 s; for the 156 cells from animal 2, the median of estimated time constant was 2.54 s with interquartile range from 0.54 to 4.39 s; for the 99 cells from animal 3, the median of estimated peak locations was 1.50 s with interquartile range from 0.40 to 3.63 s. In fact, the estimated *τ* for some cells reached the far end of the parameter search boundary (0.01-10 s), suggesting the time constants for those cells would include values beyond 10 s. We eliminated cells with *τ >* 9*s* (15/331 cells were excluded) when evaluating the correlation between *µ* and *τ* with Bayesian Kendall’s test. For cells where an accurate *τ* value can be measured, we found evidence in favor of the null hypothesis (Kandall’s *τ* = 0.07, *BF*_01_ = 2.5), indicating that there was no systematic relationship between *µ* and *τ* for past cells.

### The peaks of future cells cluster closely near the end of the interval

Similarly to the peak location of the past cells, the estimated peak locations for future cells (right side of Fig. 4b) were distributed over a narrow range of values shortly before the end of the interval (median = -0.02 s, interquartile range = 0 to -0.13 s, 90_*th*_ percentile = -0.5 s). 351/391 future cells reached peak firing within -0.5 s of interval length *T* . This pattern was true within each animal: for the 67 cells from animal 1, the median of estimated peak locations is -0.19 s with 90_*th*_ percentile = -0.62 s; for the 194 cells from animal 2, the median of estimated peak locations is -0.15 s with 90_*th*_ percentile = -0.66 s; for the 130 cells from animal 3, the median of estimated peak locations is -0.08 s with 90_*th*_ percentile = -0.29 s. This provides strong evidence against the hypothesis that time remaining before the end of the interval is coded by a group of sequentially firing cells tile the interval.

### The time constants of future cells were distributed over a broad range

Just like past cells, we observed a wide range of time constants *τ* for future cells: the median of this group was 1.03 s with interquartile range between 0.26 s and 3.00 s and the 90_*th*_ percentile was 5.39 s (See Fig. 4b). This pattern was true within each animal: for the 67 cells from animal 1, the median of the estimated time constant was 0.37 s with interquartile range from 0.17 to 1.83 s; for the 194 cells from animal 2, the median of estimated time constant was 0.80 s with interquartile range from 0.25 to 2.74 s; for the 130 cells from animal 3, the median of estimated peak locations was 2.06 s with interquartile range from 0.36s to 3.38 s. Similarly to past cells, there were a few future cells with time constant *τ* near the far end of the boundary for the future cells (10/391 cells were excluded), and were eliminated from the subsequent Bayesian Kendall’s correlation test. We found no evidence supporting a correlation between the *µ* and *τ* using the Bayesian Kendall test (Kandall’s *τ* = 0.06, *BF*_01_ = 3.13) for cells where an accurate *τ* value can be measured.

### Both time elapsed and time remaining show continuous time constants

It is of tremendous theoretical importance to know if the time constants of past cells and future cells are continuous or if they have a characteristic time scale. Visual inspection of the time constants for both past cells and future cells suggests that time constants are distributed smoothly over the parameter space (Fig. 4b). To formally test this observation, we fit the observed time constants with two types of commonly applied distributions: normal distribution and power-law distribution. The normal distribution is characterized by its expected value *µ* plus noise *σ* distributed symmetrically around the expected value. While the distribution is defined from *−∞* to *∞*, the probability of observing a value that is two factors outside of the typical value is extremely rare (<5%). Power-law distribution assumes the probability of observing *x* follows *p*(*x*) *∝ x*^*−α*^, suggesting that there is a high concentration of small values while larger values become less frequent monotonically. The distribution is characterized by its long tail where the probability of an atypical observation, no matter how far away from the concentrated small values, never asymptotes to zero. The size of the tail is determined by *α* where a smaller *α* indicates a larger portion of the observations falls under the tail. Because the long tail will continuously add more area under the density function for those less frequent but still possible (in the statistical sense) observations, power-law is only defined as a distribution between a minimum value and a maximum value for *α <* 2. In our case, a power-law distributed time constant means larger time constants gradually and *continuously* become less frequent without a clear cut-off point or distinct clusters within the bounds. Therefore one could not separate power-law distribution with a characteristic value plus noise like the normal distribution.

The time constants from the past cells and future cells were fitted separately. The fit was evaluated based on Widely Applicable Information Criterion (WAIC, Watanabe & Opper, 2010). We found that the power-law distribution provided a significantly better fit than the single normal distribution for both the past cells (WAIC for power law = 1179, WAIC for normal distribution = 1533) and the future cells (WAIC for power law = 1100, WAIC for normal distribution = 1768). We further tested if time constants should be quantified with a mixture of *two* normal distributions and found the fit improved marginally from the fit of a single normal distribution, but was still significantly lower than the power law fit for both the past cells (WAIC for mixed normal distributions = 1387) and the future cells (WAIC for mixed normal distributions = 1430). This is striking as the mixed normal distribution has five free parameters while the power law distribution only has one free parameter, the shape parameter *α*. These results provided strong quantitative evidence against the hypothesis that there exists one or few typical time scales in the population.

When the power-law exponent *α* = 1, the density function becomes *p*(*x*) *∝ x*^*−*1^, where *x* are distributed according to log scales. We found the mean of the posterior of the exponent for past cells was 0.84 with a 95% credible interval [0.75, 0.92]. The mean of the posterior of exponent for future cells was 1.07 with 95% credible interval [0.99, 1.14]. Both of the posteriors are close to 1, suggesting that the time constants were distributed *close* to logarithmic compression. When plotted against a log scale, logarithmically distributed observations would form a straight line (Cao et al., 2022). We plot the cumulative density function (CDF) on a log scale for the past cells (Fig. 4d) and the future cells (Fig. 4e) and found the corresponding curves were close to being straight, consistent with our power-law exponent estimations.

It is worth noting that the present study is not well suited to estimate the precise value of *α*. As demonstrated in Supplemental Section 3, a large portion of the population rescale the temporal receptive field with the length of reproduced interval. While our model controlled for the rescaling with additional parameter, it is possible that there is residual trade-off between the estimated *τ* and the *ρ*, especially for small *τ* . An experiment with a fixed reproduction interval would allow for a more accurate estimation of *α*.

## Discussion

We analyzed previously published recordings from mPFC during an interval reproduction task (Henke et al., 2021) using hierarchical Bayesian models to estimate the temporal receptive fields of individual cells and the populations. Consistent with previous research on interval timing (Narayanan, 2016; Merchant & Averbeck, 2017), we observed cells preferentially reach peak firing either when the animal started running, denoted as past cells, or when the animal was about to stop, denoted as future cells. Consistent with previous work (Kim et al., 2013), we saw strong evidence against a time-cell-like sequential temporal representation in mPFC cells in these data. We observed exponentially decaying temporal receptive fields for past cells and exponentially ramping temporal receptive fields for future cells. Critically, both populations showed a continuous distribution of time constants over a wide range. The firing pattern of the past cells resembled temporal context cells from entorhinal cortex (Tsao et al., 2018; Bright et al., 2020); future cells behave like a mirror image of the past cells. Future cells observed here also resembles neurons in the frontal motor cortex during the waiting period before a planned licking (Affan et al., 2024). The pattern of firing of temporal context cells resembles the real Laplace transform of the time since the interval began (see also Zuo et al., 2023). Whereas the population of past cells codes for the Laplace transform of the time since a past event, the population of future cells codes for the Laplace transform of the time until a *future* event. Together the two groups form a continuous timeline encoded in the Laplace domain, as predicted by theory (Howard, Esfahani, Le, & Sederberg, 2023).

There are several limitations to the present study. First, the present study is not well-suited to distinguish whether the animal was planning for timed future actions or replaying the memory of the stored interval during the measurement phase. These variables are highly correlated in this study. An experiment where the reproduced interval could be dissociated from the trained interval in memory, perhaps by instructing animals to produce not the interval *T*, but some factor of the interval, which would allow these two variables to be dissociated (Remington, Narain, et al., 2018). Second, the current study did not make any use of the data from the measurement phase in which the animal observed the to-be-reproduced duration for that trial. The Henke et al. (2021) paper concluded that the correlation between population vectors during the measurement and reproduction phase was rather low. Still, it is possible that including the measurement phase would produce a more complex story than considering the reproduction phase alone, as was done here.

### Future time cells can be constructed from future temporal context cells

While we observed a Laplace domain timeline in the present study, it is entirely possible, expected even, that other brain regions maintain sequentially-coded information about the time of future events, the “future time cells”. Zhou et al. (2020) observed highly sequential dynamics in the dorsolateral striatum during mice performing a two-interval timing task and argued that such dynamic provided better timing information than alternative neural dynamics including ramping. Recording from the cerebellum, Wagner, Kim, Savall, Schnitzer, and Luo (2017) observed sequentially activated cells tile time until an anticipated outcome. In hippocampus and related regions, neuroscientists have found sequential codes that appear to tile distance in physical space leading to a goal location (e.g., Gauthier & Tank, 2018; Sarel, Finkelstein, Las, & Ulanovsky, 2017).

Future cells observed here could be used directly to construct future time cells, cells that fire in sequence coding for the time until a planned event. Work on temporal coding of past events provides a strong analogy for this. It has been shown that temporal context cells, exponentially decaying cells with a continuous spectrum of time constants, can be used to generate time cells, sequentially activated cells with a continuous spectrum of peak times (Shankar & Howard, 2010; Howard et al., 2014; Rolls & Mills, 2019). Temporal context cells with a continuous spectrum of time constants can be identified with the real Laplace transform of the past. In contrast, time cells provide a blurry estimate of the past itself with different time cells tiling the time axis. Thus, the operation taking temporal context cells to time cells can be identified with the inverse Laplace transform (Howard & Hasselmo, 2020). In fact center-surround feedforward connections are sufficient to describe this operation (Shankar & Howard, 2012; Liu, Tiganj, Hasselmo, & Howard, 2019). The present study observed Laplace transform of the future—the time until a planned event. Thus, whatever neural circuit takes temporal context cells to time cells when applied to the future cells observed here would generate future time cells.

### Implications of a continuous distribution of time constants

The continuous spectra of time constants observed for both past cells and future cells are difficult to reconcile with a low-rank RNN. The dynamics defined by the recurrent matrix can have no more time constants than the recurrent matrix has unique eigenvalues. By design, a low-rank RNN has a small number of time constants. The linear dimensionality of a population with a continuous distribution of time constants is in principle unbounded. In practice, the duration of the interval studied places a practical limit on the number of dimensions that is still above what is typically proposed in low-rank RNNs.

Consider the spanning dimension for a population of past cells measured from the moment after the population peaks out to time *t*. The spanning dimension to time *t* should go like 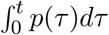 where *p*(*τ*) is the probability of observing a time constant *τ* . If *p*(*τ*) were uniform—which is clearly falsified by the results in the current paper—the spanning dimension would go up linearly with *t*. If the time constants were distributed on a logarithmic scale *p*(*τ*) *∼ τ* ^*−* 1^, the spanning dimension would increase like log *t*, a decelerating function that grows without bound. Similar results would be observed for the population of future cells if the integral was started at the end of the delay rather than the beginning. Studying neural data from working memory experiments in monkeys, Cueva et al. (2020) observed that the dimensionality spanned by neural population indeed increased sublinearly with recording time.

### Continuous attractor networks for exponential ramping

Several researchers have modeled temporal coding in the brain using RNNs (Laje & Buonomano, 2013; Remington, Egger, et al., 2018; Beiran et al., 2023). But the dynamics observed here—in particular the population of future cells that grow exponentially with a continuous spectrum of time constants—present a serious challenge for recurrent networks. First, note that it is straightforward to write an RNN that gives out a population of exponentially decaying or ramping cells as a solution. For instance, a population of past cells could be simply constructed from a recurrent connection with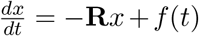, where *x*(*t*) is the recurrent state and *f* (*t*) is the external input at time *t*. If an input is given to all units in *x* at time zero, then one obtains *x*(*s, t*) = *e*^*−st*^. If **R** is a diagonal matrix with the appropriate values of *s* along the diagonal. However, this solution does not work for ramping neurons. Although we can write down a solution by initializing the network at *t* = 0 with *f* (*t* = 0) = *e*^*−sT*^ and changing the sign of **R**, this solution would be extremely unstable. Even tiny amounts of noise in the initial state would be amplified by exponential growth. After a short period of time, the pattern of activation across different cells would no longer be reliable or coherent.

Happily, if we restrict our attention to networks with a logarithmic distribution of time constants (Cao et al., 2022), the problem is much more constrained and can be solved without a conventional RNN. Figure 5 illustrates the basic insight: If the values of *s* in the population are distributed logarithmically, then a plot of 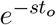 as a function of cell number appears as an edge for any particular value of *t* (within bounds specified by the minimum and maximum values of *s*). Comparing 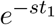 to 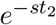 on this graph merely shifts the location of the edge. This suggests that one could build a continuous attractor neural circuit that supports a stable edge to implement these equations. In this circuit, the difference between ramping neurons and decaying neurons would simply be the direction and rate at which the edge moves across the population as time progresses (for more detail see Howard et al., 2023). Ensuring that the individual neurons follow an exponential time course requires careful construction of the connectivity between neurons in the attractor network. For all choices of connectivity, the time evolution will be scale-invariant—expressible as *t/τ*_*i*_, where *τ*_*i*_ is specific to neuron *i*—as long as the time constants are logarithmically-compressed and the edge translates as a function of time.

**Figure 5.**
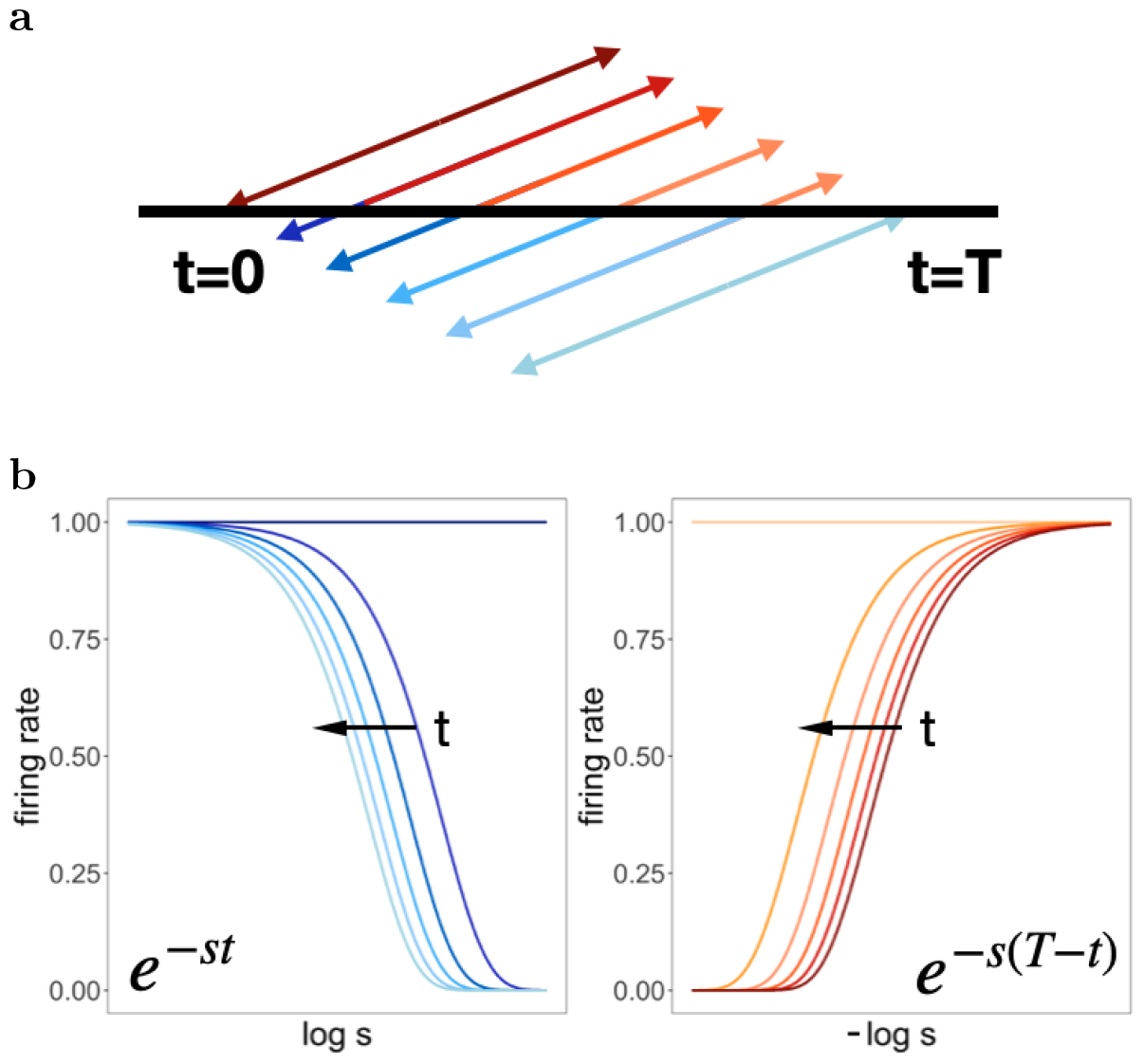
Continuous attractor network for exponential decay and ramp with logarithmically-spaced time constants. **a**. The interval begins at *t* = 0 and ends at *t* = *T* . During this period the memory for the beginning of the interval recedes from the present towards the past and the expectation of the end of the interval approaches from the future closer to the present. **b**. The plots on the left show *e*^*−st*^ for several choices of *t* as a function of log *s*. The plots on the right show function *e*^*−s*^(^*T −t*^) for several choices of *t* as a function of *−*log *s*. The value of *t* is communicated by the shade of the line, corresponding to the moments in **a**. Note that the shape of the pattern of activity over neurons does not change, for either the past or the future. Thus a continuous attractor network would be able to account for these results. This is only possible if time constants are evenly distributed on a log scale. Although the difference between the values of *t* is the same, the distances between adjacent edges are not the same. This is because *s* is on a logarithmic scale. Note that whereas the distance between successive lines becomes progressively smaller as the edge recedes into the past, the distance between successive lines becomes progressively larger as the future approaches.

It is worth noting that this view of time is not unlike “thermometer variables” observed in the sensory organs. For instance, think about loudness thresholds in auditory-nerve fibers (Sachs, Winslow, & Sokolowski, 1989). Individual fibers modulate their firing over a small dynamic range of loudness around some threshold value, with firing saturating at higher values. Different fibers have different thresholds and the dynamic range of individual fibers is much less than the dynamic range over which the animal can distinguish loudness. This means, for any particular loudness, the population can be divided into fibers that are above their threshold and neurons that are below their threshold. Sorting the fibers on their thresholds we would observe an edge. As loudness changes, the “location” of this edge changes. This neural motif—of a continuous attractor network with an edge—could be used for any situation where a neural population needs to represent an individual scalar value. In this study, the scalar value represented by the past cells is time since the beginning of the interval; the scalar value represented by the future cells is time until a planned movement.

## Materials and Methods

Because we took the recording data from Henke et al. (2021), the specific details about the data acquisition can be found in the original publication. In the section below we describe the computational analysis applied to the data in detail.

### Data analysis

The goal of these analyses is to identify parameters that best describe the properties of time fields of temporally responsive cells at both the trial and population levels. First, we identify and characterize temporally responsive cells using previously established methods (Tiganj et al., 2018; Bright et al., 2020). Then we apply Bayesian models to capture the temporal properties across trials and of the population.

#### Selecting temporally responsive cells as input to hierarchical Bayesian model

In order to initially identify temporally responsive cells, spikes were first analyzed in Matlab 2016a using a custom maximum likelihood script. This approach has previously been used to identify both time cells and temporal context cells (Tiganj et al., 2018; Bright et al., 2020). Fits were restricted to spikes that occurred during the reproduction phase. We fit the spike trains to five models. The first model was a constant firing model, which indicates a lack of temporal responsiveness. The constant firing model,

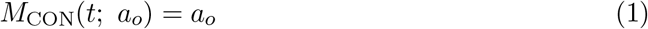

contained a single parameter *a*_*o*_, allowed to range between 10^*−*7^ and 1, which predicted a constant probability of firing a spike at time *t*.

We then considered an exponentially modified Gaussian model, which describes the firing field as a convolution of an exponential distribution and a Gaussian distribution. This model was fit to both time elapsed since the start of the reproduction phase and time remaining until the end of the reproduction phase. The exponentially modified Gaussian model for time since the start of the reproduction phase,

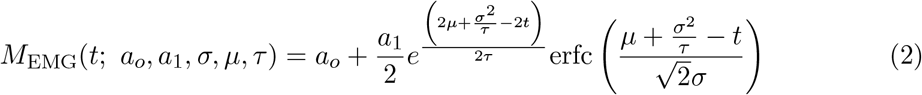

contained the parameters *t*, which tracked time since the start of the reproduction interval, *a*_*o*_ (allowed to range between 10^*−*7^ and 0.5) describes the baseline firing rate, *a*_1_ (allowed to range between 10^*−*7^ and 0.5) describes the amplitude of the temporal field, *µ* (allowed to range between 0 and the average midpoint across reproduced intervals for a given cell) describes the temporal receptive field peak location of the cell, *σ* (allowed to range between 0.01 and 0.5) describes the variability of the peak, *τ* (allowed to range between 0.1 and 16) describes the time constant of the exponential decay in firing, and *ecrf* is the complementary error function.

In addition, we also considered an exponentially modified Gaussian model that modulated the exponential time constant by the length of the reproduced interval. This reproduction length modulated model replaced *τ* with *τ*_*i*_, such that 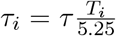 where *τ*_*i*_ linearly rescale with reproduced interval length *T*_*i*_. Parameters fit by the model spanned the same ranges as in the prior exponentially modified Gaussian model.

We also fit the exponentially modified Gaussian model for time until the end of the reproduction phase, replacing *t* with *T − t*, with models including both with *τ* and *τ*_*i*_. All parameters fit by the maximum likelihood models for the end of the delay spanned the same ranges as the parameters in prior models.

We evaluated the models for each neuron using a likelihood ratio test and counted the number of neurons that were better fit by an ex-Gaussian model at the 0.05 level, Bonferonni-corrected by the total number of neurons. We further required that both even and odd trials for a neuron were significantly fit by the same model, and that those fits had a Pearson’s correlation coefficient greater than 0.4. In the event that multiple ex-Gaussian models met these requirements for a given cell, we selected the model with the best log-likelihood. Therefore the temporal responsive cells are further categorized into cells best fit with an ex-Gaussian model with time elapsed (*t*, the “past cells”), and cells best fit with an ex-Gaussian model with time remaining (*T − t*, “future cells”).

#### Hierarchical Bayesian models describe temporal receptive field across trials

We fit hierarchical Bayesian models inspired by the results from the maximum likelihood methods, with additional parameters estimated at the trial level so cells that code for absolute time and cells that rescale with different reproduced intervals can be captured with the same model. A graphic diagram of the Bayesian model can be found in Fig. 6

**Figure 6.**
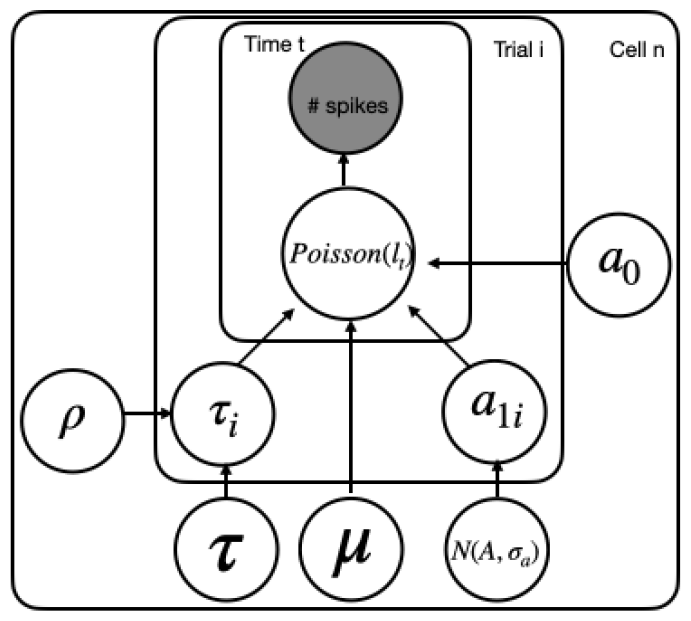
Illustration of the hierarchical Bayesian model. Each node represents a variable in the model; the filled node represents the observed number of spikes in each time bin (10ms) and open nodes represent latent variables. Arrows represent relationships between variables and plates indicate whether the variable is estimated at the trial level, or cell level.

For cells categorized as past cells by the maximum likelihood methods, we assume a Poisson process binned at 10 ms is applied to evaluate the latent model *M*_*EMG*_. The model is largely the same as Eq. 2 except that both time constant *τ* parameter and amplitude of time field *a*1 are estimated for each trial i:

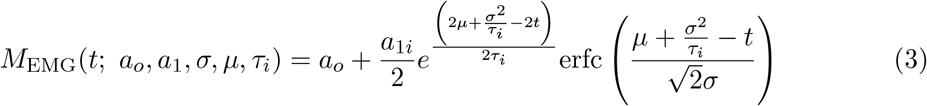

The trial level time constant *τ*_*i*_ is modified according to the time constant *τ* (allowed to range between 0.1 to 10) at 5.25 s with an additional free parameter *ρ* and the interval length *T*_*i*_:

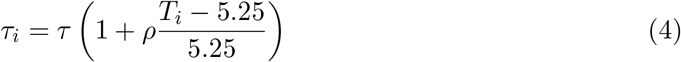

where *ρ* quantifies how much and in which directions time constant *τ*_*i*_ changes relative to the change of the produced interval length from the baseline estimation *τ* when the interval is 5.25 s. A positive *ρ* indicates the cell increases its time constant for longer intervals while a negative *ρ* indicates the opposite. It is obvious that a *ρ* close to zero means the cell code for absolute time and does not change its time constant for different intervals. Another value of interest is when *ρ* = 1. In this case the time constant *τ*_*i*_ linearly rescales with the interval: for example, if the interval is doubled (*T*_*i*_ = 10.5*s*), the corresponding *τ*_*i*_ would also be twice the *τ* . When the absolute value of *ρ* is smaller than 1, it means *τ*_*i*_ changes relatively less compared to the change in the interval length. *ρ* are allowed to take any value as far as the resulted *τ*_*i*_ from Eq. 4 is positive for the temporal receptive field estimation.

Another trial-level parameter *a*1_*i*_ is allowed to vary freely across trials. The only constraint was applied indirectly through marginal probability with the assumption that *a*1_*i*_ is sampled from a normal distribution for a given cell. In addition, *σ* from the ex-Gaussian field is fixed to 0.1 for all cells because the parameter resulted consistently in small values from the maximum-likelihood method. For cells categorized as future cells, we applied the same model as Eq.3 except replace time elapsed *t* with time remaining *T*_*i*_*− t* for each trial. For the past cells the peak time *µ* is allowed to range between 0 to 3 s from the start of each trial; for the future cells the peak time *µ* is allowed to range between 0 to 3 s to the end of each trial.

The posterior distributions of estimated parameters were generated through the rStan package (Stan Development Team, 2022) with 4 independent Markov-Chain-Monte-Carlo (MCMC) chains (`5200 warm-up iterations` and `50 post warm-up samples` in each of the MCMC chains). This procedure returns posteriors of the likelihood of the data as well as posteriors for the parameters of the model.

#### Evaluating the time constants population with Bayesian models

We compared three models (power-law distribution, single normal distribution, and a mixture of two normal distributions) to describe the time constants population for the past cellsthe future cellsseparately, *after* the hierarchical Bayesian models described above were applied to quantify the temporal receptive field for each cell. The distribution parameters were estimated through rStan package with 8 indepdent MCMC chains(`4000 warm-up iterations` and `4000 post warm-up samples` in each of the MCMC chains).

#### Power-law distribution

The probability density function of Power law distribution can be derived within the range [min, max] of time constants *τ* :

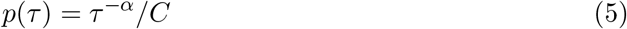

where

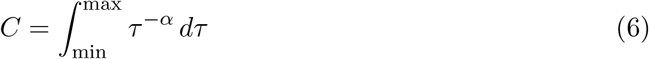

is the area under the density function over [min, max].

When *α* ≠ 1, the result of the integration is:

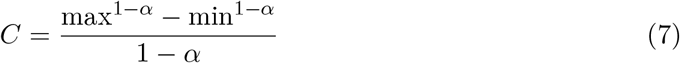

When *α* = 1, the result is:

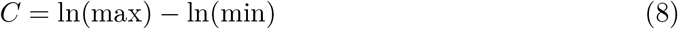

When fitting the Power-law model, we adopt a straightforward approach by setting the upper and lower bound of the power-law to be the same as the parameter space of *τ* .

#### Normal distribution

In the second model, we quantify the time constants with a single normal distribution:

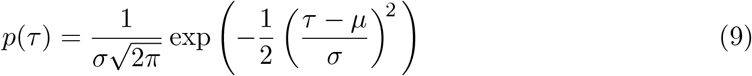

where *µ* is the estimated stereotypical time constant for the population.

#### Mixture of two normal distributions

The third model we tested is a mixture model of two normal distributions with the probability of *τ* sampled from the first normal distribution to be *λ*_1_ and the probability of the second normal distribution to be 1 *− λ*_1_:

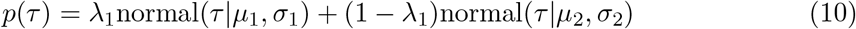

## Supplementary Materials

### S1: Non-temporal cells show similar patterns to the temporal responsive cells

In Fig. S1 we report the temporal properties across different stimulus intervals for cells where the maximum likelihood methods failed to identify a temporal receptive field (non-temporal cells, 1044/1766). As we did with temporal responsive cells, we separated the non-temporal cells into two groups: one group where hypothetical temporal receptive fields from the start of the interval provide better fits (analogous to the past cells, 489/1044), and the other group where hypothetical temporal receptive fields from the end of the interval provide better fits (analogous to the future cells, 477/1044). 79 cells were eliminated for missing data in at least one stimulus interval. Despite lacking statistically significant temporal receptive fields, the non-temporal cells visually resemble the firing pattern of the temporal cells in Fig.2. Therefore we performed the same hierarchical Bayesian models on the non-temporal cells as the temporal responsive cells in the main article.

### Non-temporal cells are characterized with a broad range of time constants

Fig. S2d summarizes the estimated peaks and time constants of the temporal receptive fields of non-temporal cells. Although the cells are plotted as separated groups, we report the estimated peaks *µ* and time constants *τ* together due to the high similarity between the two groups.

Like the temporal responsive cells in the main text, the time peaks *µ* tend to cluster around the start and the end of the interval (median = 0.06 s, interquartile range = 0.01 to 0.38 s, 90_*th*_ percentile = 1.08 s). This is a broader range than the temporal responsive cells, but still rules out sequential tiling possibility.

Similarly, the time constants *τ* for the non-temporal cells also resemble what we observed for the temporal responsive cells. Like the temporal responsive cells, the estimated time constants *τ* cover a broad range of values (median = 3.57 s, interquartile range = 0.53 to 9.04 s, 90_*th*_ percentile = 9.88 s). There is a significant portion of cells with *τ* at the boundary of parameter search space (>9 s), meaning that the firing rate changes very little during the interval for those cells. It is likely that these cells maintain a constant firing rate throughout the interval. After eliminating those cells (246/966), we plotted the cumulative density functions of the non-temporal cells on a log scale in Fig. S1c and d. The resulting density functions highly resemble the ones of temporal responsive cells in Fig. 4d and e.

**Figure S1.**
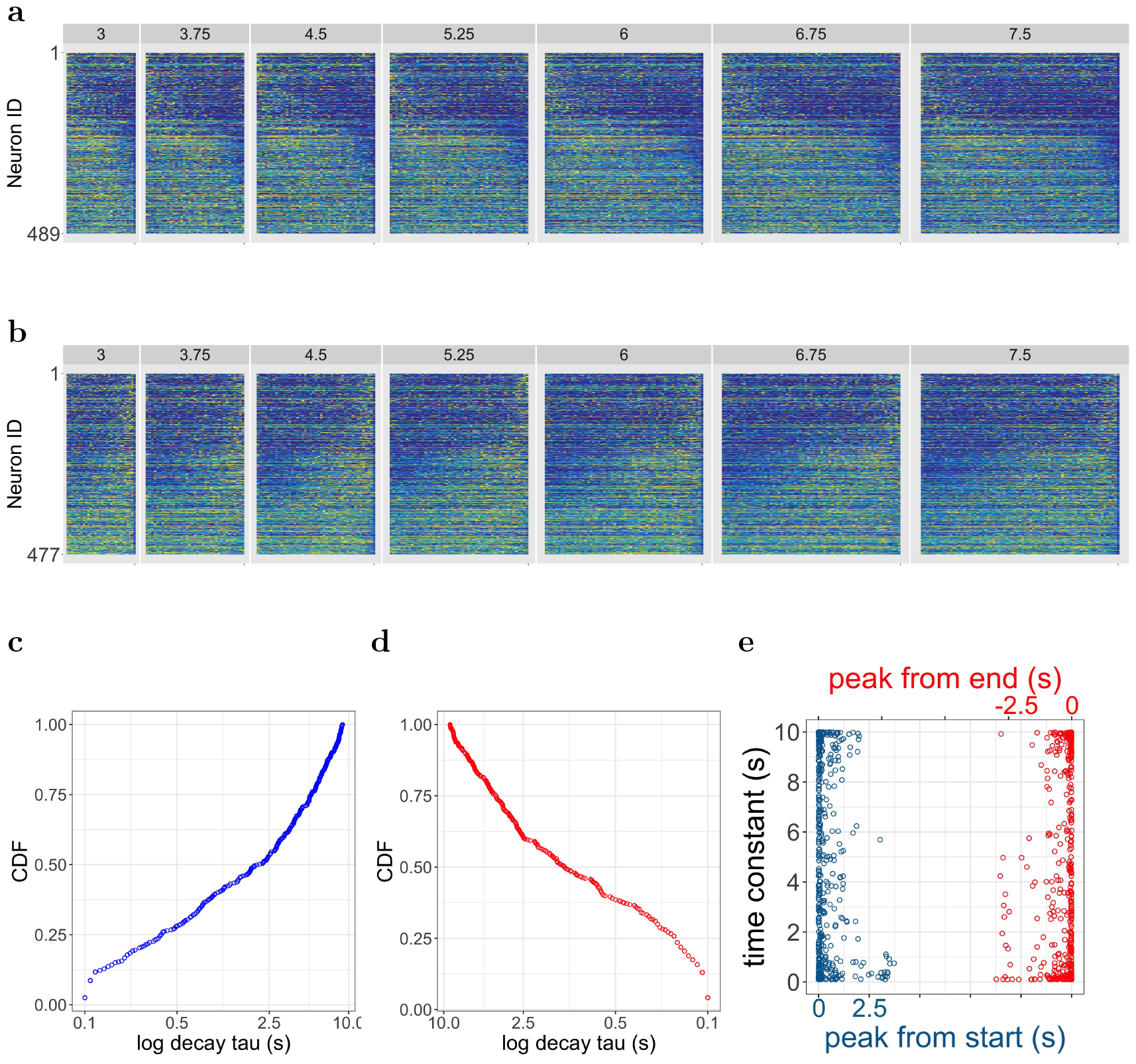
Non-temporal cells resemble a noise version of the temporal responsive cells. **a-b**. Heatmaps of non-temporal cells during reproduction for all delay lengths. Heatmaps plotted in the same way as Fig. 2. Like the temporal responsive cells reported in the main text, the non-temporal cells can be divided into two groups: one with firing patterns better aligned with the start of the interval (**a**) and the other with firing patterns better aligned with the end of the interval (**b**). **c-d**. Cumulative density functions of the estimated time constants for non-temporal cells on a log scale. The plots follow the same format as Fig. 4d and e. Cells with estimated time constants at the parameter space boundary (>10) are omitted in the plot. **e**. Estimated time constants as a function of estimated peak locations. Plotted in the same way as Fig. 4b.

### S2:Additional example cells

**Figure S2.**
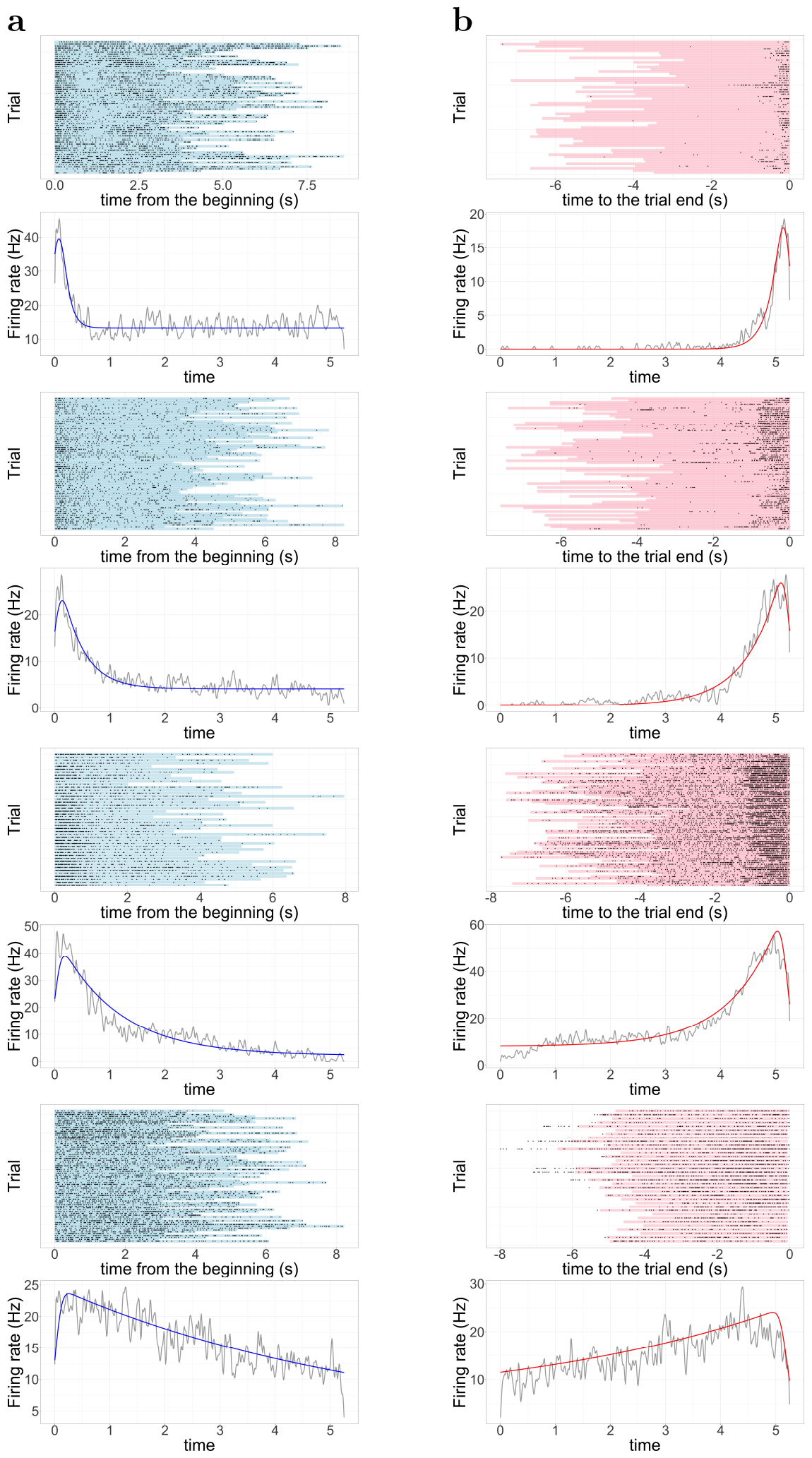
Additional temporal cell fits. Format is the same as Figure 2.

**Figure S3.**
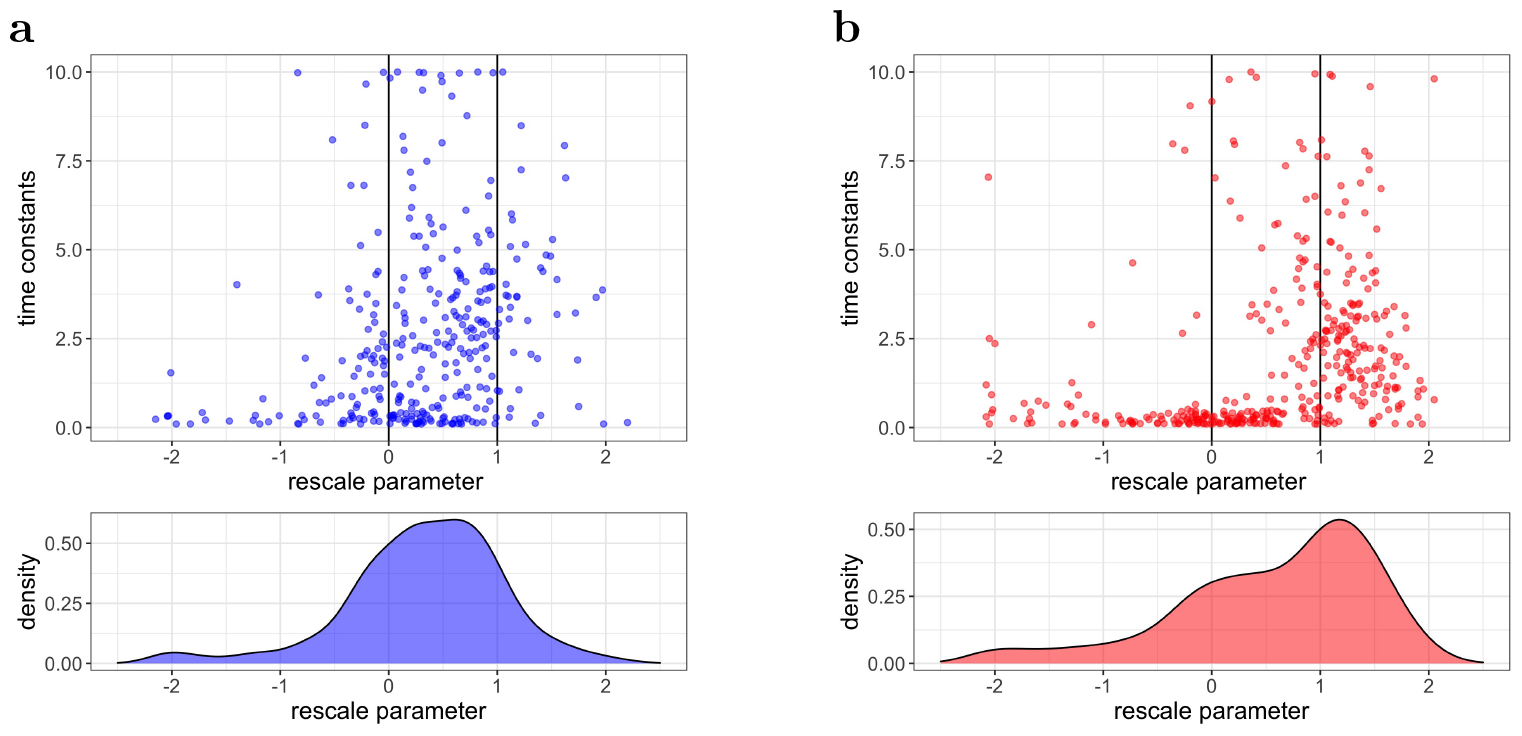
The future cells are more likely to increase time constants for longer intervals than the past cells. **a**. Rescale parameter *ρ* as a function of time constant *τ* (top) and the density distribution (bottom)for the past cells. The majority of cells are distributed between coding for “absolute time” (*ρ* = 0) and “linearly rescale” (*ρ* = 1). **b**. The same as **a**. but for the future cells. There are significantly more cells linearly rescale their temporal receptive field (*ρ* = 1) among the future cells than the past cells.

### S3: The future cells are more likely to rescale with interval length than the past cells

In the Results section of the paper, we report the peak location parameter *µ* and the time constant parameter *τ* that quantifies the temporal receptive field for each cell. Below we report the rescale parameter *ρ* that explicitly quantifies how the temporal receptive field changes according to Eq. 4 in the Methods section. Specifically, when *ρ* is around 0, the temporal receptive field changes very little for different intervals, and when *ρ* is around 1, the temporal receptive field linearly rescales to the interval length. Fig. S3 summarizes the rescale parameter *ρ* for the past cells (left) and the future cells (right). Cells in both groups show a spectrum of sensitivities to the interval length, while a larger portion of the future cells rescales linearly with the interval length.

In the top panels of Fig. S3 we plot *ρ* as a function of the corresponding time constant *τ* for the past cells (a) and the future cells (b). The estimation of *ρ* for cells with small time constants is rather noisy, and we believe the observations with large absolute values of *ρ* (>1.5) are mostly the results of such noise. In the bottom panels of Fig. S3 we plot the distribution of *ρ* for the past cells (a) and the future cells (b). For the past cells, the majority of cells mildly increase the trial-level time constant with the rescale parameter *ρ* placed between 0 to 1 (median = 0.37, interquartile range = -0.07 to 0.81, 90_*th*_ percentile = 1.12). By contrast, the distribution of the rescale parameter for the future cells peaks around 1 (median = 0.8, interquartile range = -0.03 to 1.26, 90_*th*_ percentile = 1.52). The difference between the distribution of the past cells *ρ* and the distribution of the future cells *ρ* is significant based on KS test (D=0.29, p-value <0.001). This finding was consistent with report in Henke et al., 2021 using none parametric methods.

## Acknowledgments

The authors gratefully acknowledge support from R01MH132171 and a Google-AI Faculty Research Award.

## Footnotes

1 on the manifold defined by the low-rank recurrence matrix

